# Mixture coding and segmentation in the anterior piriform cortex

**DOI:** 10.1101/2019.12.26.888693

**Authors:** Sapir Penker, Tamar Licht, Katharina T. Hofer, Dan Rokni

**Affiliations:** Department of Neurobiology, School of Medicine and IMRIC, The Hebrew University, Jerusalem, Israel

## Abstract

Coding of odorous stimuli has been mostly studied using single isolated stimuli. However, a single sniff of air in a natural environment is likely to introduce airborne chemicals emitted by multiple objects into the nose. The olfactory system is therefore faced with the task of segmenting odor mixtures to identify objects in the presence of rich and often unpredictable backgrounds. The piriform cortex is thought to be the site of object recognition and scene segmentation, yet the nature of its responses to odorant mixtures is largely unknown. In this study, we asked two related questions. 1) How are mixtures represented in the piriform cortex? And 2) Can the identity of individual mixture components be read out from mixture representations in the piriform cortex? To answer these questions, we recorded single unit activity in the piriform cortex of naïve mice while sequentially presenting single odorants and their mixtures. We find that a normalization model explains mixture responses well, both at the single neuron, and at the population level. Additionally, we show that mixture components can be identified from piriform cortical activity by pooling responses of a small population of neurons - in many cases a single neuron is sufficient. These results indicate that piriform cortical representations are well suited to perform figure-background segmentation without the need for learning.

## Introduction

The odorants emitted by different objects in the environment mix in the air before reaching the nose. Each of these objects in itself will typically emit tens to thousands of odorants that become its olfactory signature. Natural inputs into the olfactory system are therefore rich odorant mixtures that require segmentation in order for useful information to be extracted. The difficulty of mixture segmentation arises from the overlapping representations of odorants by olfactory sensory neurons (Malnic et al., 1999; Rubin and Katz, 1999; Araneda et al., 2000; Kajiya et al., 2001; Abaffy et al., 2006; Grosmaitre et al., 2009; Soucy et al., 2009). Similar to the auditory system (and unlike the visual system), a single sensory neuron may be simultaneously activated by multiple odorants (Brungart et al., 2001; McDermott, 2009). Behavioral testing has indeed shown that increased overlap in odorant representations is related to increased difficulty of scene segmentation (Rokni et al., 2014).

The mechanisms for scene segmentation are not well understood. Piriform cortex, being the first brain station that combines inputs from multiple receptor channels, is a primary candidate for performing this task (Davison and Ehlers, 2011; Ghosh et al., 2011; Sosulski et al., 2011; Haddad et al., 2013). Several factors have been suggested to contribute. First, if objects are dispersed in space, plume dynamics may provide temporal separation between objects (Hopfield, 1995; Brody and Hopfield, 2003; Wilson, 2003; Kadohisa and Wilson, 2006a; Linster et al., 2007; Szyszka et al., 2012; Vinograd et al., 2017). Second, readout of mixture components can be achieved by properly combining inputs form multiple receptor channels (Mathis et al., 2016; Singh et al., 2019). And third, feedback projections within the olfactory system may provide a flexible readout for specific mixture components (Li and Hertz, 2000; Grabska-Barwińska et al., 2017; Singh et al., 2019).

Currently a limiting factor in understanding mixture segmentation, is our limited knowledge about mixture coding. Understanding how odor mixtures are represented in the piriform cortex is a prerequisite for understanding mixture segmentation. Ultimately we would want to be able to predict responses to mixtures based on the responses to single components. Several studies have reported sublinear mixture responses in olfactory cortex, thereby limiting the possible response space, however they have not provided models to predict mixture responses (Lei et al., 2006; Yoshida and Mori, 2007; Stettler and Axel, 2009). Sublinearlity of mixture responses is probably inherited to some extent from the olfactory epithelium (Kurahashi et al., 1994; Duchamp-Viret et al., 2003; Oka et al., 2004; Takeuchi et al., 2009; Xu et al., 2020; Zak et al., 2020), as well as from the olfactory bulb where cross odorant inhibition may contribute (Yokoi et al., 1995; Urban, 2002; Aungst et al., 2003; McGann et al., 2005; Arevian et al., 2008; Fantana et al., 2008). The piriform cortex integrates these non-linear odor representations from the olfactory bulb and utilizes local recurrent circuitry to generate representations that presumably support segmentation (Poo and Isaacson, 2009, 2011; Franks et al., 2011; Miura et al., 2012; Suzuki and Bekkers, 2012; Roland et al., 2017; Bolding and Franks, 2018). Importantly, piriform cortex is also expected to contribute to sublinear summation of mixture components due to local inhibitory circuits. These circuits have been shown to normalize responses to increasing concentrations of odors (Bolding and Franks, 2018; Stern et al., 2018). Whether increasing stimulus intensity by increasing concentration or by adding more odorants produces equivalent normalization is unknown.

Several questions remain unanswered about mixture representations. First, how do cortical mixture representations relate to the representations of their constituent odorants? In other words, how can one predict cortical responses to mixtures from the responses to single components? And second, how is information about the individual mixture components represented in the piriform cortex and how can it be read by down-stream regions?

To answer these questions, we systematically characterized mixture representations in the piriform cortex and analyzed their ability to convey information about individual mixture components, at both the single neuron, and population levels.

## Materials and Methods

All experimental procedures were performed using approved protocols in accordance with institutional (Hebrew University IACUC) and national guidelines.

### Data acquisition

Young adult male c57bl6 mice (10-14 weeks old, Envigo) were anesthetized (Ketamine/Medetomidine 75 and 1 mg/kg, respectively), and were restrained in a stereotaxic device (Model 940, David Kopf Instruments). The skin was removed from the scalp and a small craniotomy was made over the anterior piriform cortex (1.5 mm anterior and 2.8 mm lateral to bregma). A metal plate was attached to the skull with dental acrylic and was used to hold mice at the electrophysiological rig. Normal body temperature (37 C) was maintained with a heating pad (Harvard Apparatus). A single tungsten electrode (A-M systems, 10-12 MOhm) was lowered into the piriform cortex with a micromanipulator (Sutter Instruments). Signals from the electrode were band pass filtered (300-5000 Hz), amplified (X1000, A-M systems 1800), and sampled at 20KHz and digitized with 16-bit precision (National Instruments PCIe-6351). All analog signals were displayed and saved for offline analysis using custom-made software in LabVIEW (National Instruments). Injections of a TRITC-labeled 10Kd dextran (500 nL, Molecular probes cat#D1817, 10mg/ml) were used to verify piriform cortical targeting.

Odors were presented using a custom-made, computer-controlled, odor presentation machine that maintains constant flow and allows mixing of odors without mutual dilution (Rokni et al., 2014). The following 8 odorants were used (all from Sigma Aldrich, CAS numbers in parentheses): 1. isobutyl propionate (540-42-1), 2. 2-ethyl hexanal (123-05-7), 3. ethyl valerate (539-82-2), 4. propyl acetate (109-60-4), 5. isoamyl tiglate (41519-18-0), 6. phenethyl tiglate (55719-85-2), 7. citral (5392-40-5), 8. ethyl propionate (105-37-3). All odorants were diluted to 10% in diethyl phthalate (84-66-2), and then further diluted 8 fold in air. Odors were presented into a mask that fit the mouse’s snout at a rate of 1 l/min and were cleared from the mask by vacuum that was set to create an outward flow of 1 l/min (Figure 1A). A third port of the mask was connected to a mass flow sensor (Honeywell AWM3300) to monitor respiration. For each recording session, a set of 4 odorants was chosen (initially randomly and later fixed to odorants 1, 3, 5, and 7). All 15 combinations of these 4 odorants were then presented with randomized order (typically repeating ~30 times each). Odors were presented for 1.5 s with an inter-trial interval of 20 s.

**Figure 1.**
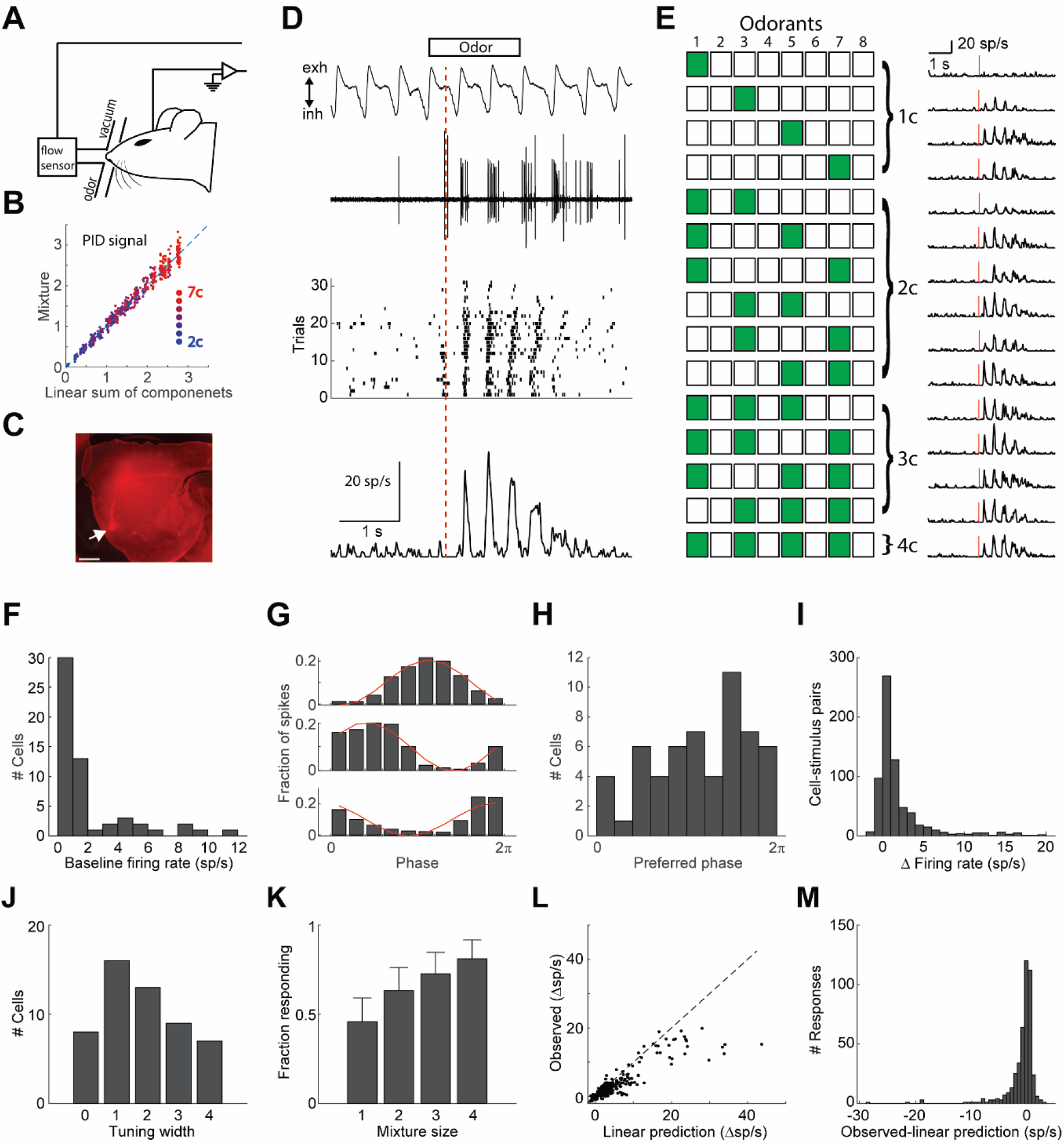
Mixture responses in piriform cortex. **A.** Schematic of experimental setup. **B.** Odor presentation calibration. The PID signal for mixtures plotted vs the linear sum of PID signals for mixture components, indicating that mixture stimuli are a linear sum of the single component stimuli. Dot color indicates mixture size. **C.** A sagittal section of a brain that was injected with a 10 kDa rhodamine at the same coordinates used for recordings. Arrow marks the injection site in the anterior piriform cortex. **D.** Example raw data (respiration and electrophysiology), raster plot, and PSTH in response to a mixture of 2 odorants (od 3&5). Red line denotes the time of PSTH alignment (first inhalation onset). **E.** An example stimulus set. All combinations of 4 odorants. Green-filled and empty squares denote odor on and off, respectively. PSTHs on the right are the responses of the same cell as in D to all stimuli. Red lines show the time of the first inhalation onset. **F.** Histogram of baseline firing rates. **G.** Fraction of spikes as a function of respiratory phase shown for 3 example cells. Red line shows a cosine fit. **H.** Histogram of preferred baseline firing phase. **I.** Histogram of change in firing rate in response to odor stimulation. **J.** Histogram of the tuning width of all cells that responded to at least one of the fifteen stimuli. Tuning width is defined as the number of single odorants that elicited a statistically significant response. **K.** Fraction of significant responses as a function of the number of odorants in the mixture. Error bars show the 95% confidence interval. **L.** Observed responses (firing rate) vs the linear sum of individual odorant responses. Responses of all cell-mixture pairs are shown. Dashed line is the unity line. **M.** Histogram of the differences between observed and linearly predicted mixture responses.

### Data analysis

Action potentials were detected offline using custom-written code in Matlab, and were then sorted using MClust (David Redish). Only units that had less than 1 in 1000 spikes to have occurred within a 3 ms refractory period were considered as single units and further analyzed. Respiratory signals were low-pass filtered with a moving average window of 250 ms. Inhalation onsets were defined as zero crossings with negative slope. All neural responses were aligned to the first inhalation onset following odor onset. PSTHs were generated with 1 ms bins and filtered with a 50 ms window Gaussian moving average. Response significance was tested by comparing spike rates during a 3 second response period to the spike rates in a 3 second period with no odor, using the Wilcoxon ranksum test with a threshold p-value of 0.01. All responses are shown as baseline-subtracted spike rates. To fit normalization models, data was fit to equation 1 using the lsqcurvefit function in Matlab. Fit quality was assessed with the coefficient of determination (R^2^).

Receiver operating characteristic (ROC) analysis was performed using the perfcurve function in Matlab. For each neuron, ROC analysis was performed separately to test the detectability of each odorant. The reported performance of each neuron is the best across the 4 odorants. The performance reported for each odorant is the best across all neurons. Shuffled controls were created by shuffling the stimulus labels for all neurons and obtaining a distribution of 224 auROC values (4 odorants X 56 neurons). Performance is reported as above control if the auROC was above all shuffled controls.

For population analysis, we generated a pseudo-population by pooling neurons from different experiments in which we used the same 4 odorants (odorants 1, 3, 5, and 7). Principal component analysis was performed after normalizing each neuron’s responses to its standard deviation. For modelled mixture responses we created modelled PSTHs for each neuron. The linear, mean, and max modelled PSTHs were created by taking the linear sum, mean, and maximum of the individual component responses, respectively. The normalization model PSTHs were created by simultaneously fitting a pair of parameters to all mixture responses of a neuron (equation 1).

Linear classifiers were realized as logistic regressions and were fit using the fitclinear function in Matlab. Classifiers were cross-validated by creating a test set that was not used for training. Each test set included one randomly selected response from each neuron to each of the 15 odor combinations. A training set that did not include the testing trials was then generated by randomly selecting trials from each neuron for each odor combination. The training set included 10,000 trials of each of the 15 stimuli for most analyses. For the analysis of temporal resolution, the training set was increased to 100,000. The reported performance for the classifiers is the mean over 100 iterations. For assessing the effect of population size on classification accuracy, we gradually removed inputs from the classifier by sequentially removing the neuron with the minimal absolute weight.

## Results

We recorded the responses of single neurons in the anterior piriform cortex of naïve anesthetized mice, to all possible combinations of 4 odorants (4 single odorants, 6 pairs, 4 triplets, 1 mix of 4, Figure 1A-E). Various sets of 4 odorants were used. Because the isolation of single-unit activity is critical for the following analysis, we recorded neural activity using single tungsten electrodes which provide a much higher signal to noise ratio than most multi-channel systems and allows better separation of single-unit spikes. We recorded from 56 well-isolated neurons of which 53 responded significantly to at least one of the 15 odor combinations. Odors were presented using a machine that allows mixing of odors without mutual dilution, into a mask that fit the mouse snout and were cleared with constant negative pressure (Figure 1A,B). The mask was connected to a mass flow sensor for continuous monitoring of respiration. Dye injections in the recording coordinates (1.5 mm anterior and 2.8 mm lateral to Bregma, 4.5mm ventral), verified targeting of anterior piriform cortex (Figure 1C). All odor responses were aligned to the first inhalation onset following odor valve opening, and typically showed a strong locking to respiration (Figure 1D,E).

The basal firing of cortical neurons was typically very low (1.9 ± 0.05 Hz, mean ± SEM, Figure 1F), and both basal firing and odor responses were typically strongly modulated by respiration (Figure 1E, G, & H). Odor responses were predominantly positive (increases in firing rate) with only 18% showing mild decreases in firing rate (Figure 1I). This however may be an underestimate of inhibitory inputs as most neurons had a basal firing rate of less than 1 spike per second, rendering our analysis less sensitive to inhibition. Of the responding neurons, most responded significantly to more than one of the individual odorants (Figure 1J). We first asked whether adding odorants to a mixture increases overall responses in the piriform cortex. Analyzing response statistics across all responsive neurons, we found that the fraction of significant responses (p<0.01) was positively correlated with the number of mixture components (r=0.98, p=0.017, Figure 1K). The simplest model for response integration is a linear model in which a neuron’s response to a mixture is equal to the sum of its responses to the mixture components. To test for linearity of odorant integration, we compared mixture responses of all neurons to the linear sum of the responses to individual mixture components. Most mixture responses were below the linear prediction (Figure 1L,M). These analyses show that in agreement with previous studies, the vast majority of piriform cortical responses to odorant mixtures increase with added mixture components and that this increase is sublinear.

We next asked whether there is any consistent shape to the relationship between the linear predictions and the observed responses. We first treated responses as the mean firing rate within a 3 seccond response period. We fitted mixture responses with a logistic function to describe response normalization, similarly to fits used to describe mixture responses in olfactory sensory neurons (Mathis et al., 2016):

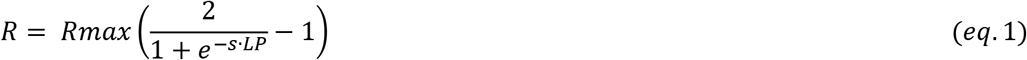

In this model, *R* is the neuron’s response to a mixture, *LP* is the linear sum of the individual component responses, *Rmax* is the neuron’s maximal response, and *s* is a parameter that sets the initial slope of the function (at LP=0). *Rmax* and *s* are the two fitted parameters. We found that this model explains mixture responses well for most neurons (Figure 2A1-A3). Some neurons however, could not be properly fit (Figure 2A4). The normalization model could account for most of the variance (R^2^ above 0.5) in 30/53 (57 %) of the cells (Figure 2B). The neurons that could not be well explained with this model (low R^2^) showed little increase in their responses as mixture components were added as assessed by the slope of a linear fit between the linear prediction and the observed responses (Figure 2C). Whether these neurons represent a separate neuronal subtype is unclear, yet the quality of the normalization fit was not correlated with spike waveform, basal firing rate or preferred respiratory phase of firing (Figure 2D-F). These results indicate that mixture responses of individual piriform neurons can often be explained by a simple normalization model.

**Figure 2.**
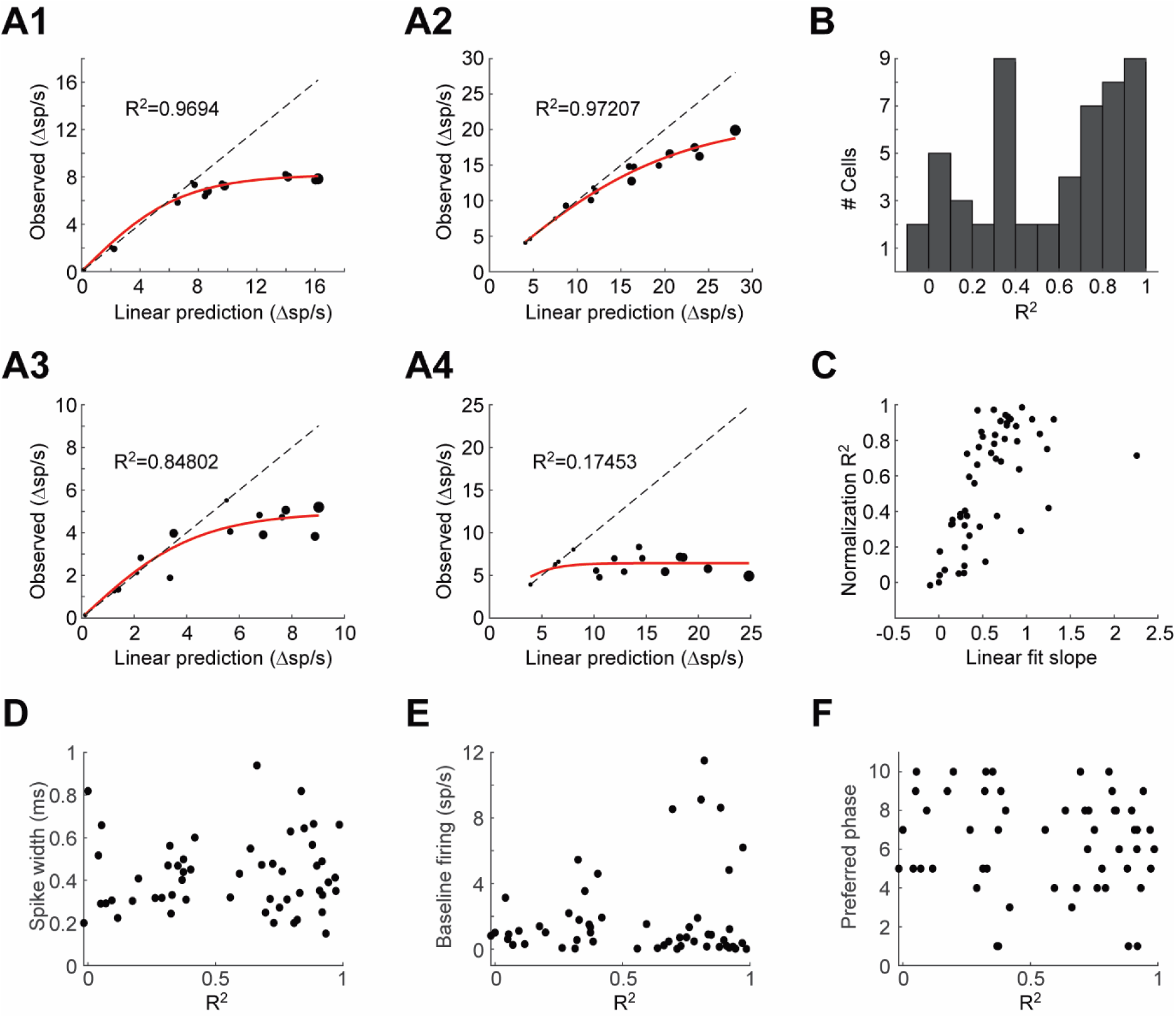
A normalization model for mixture responses of individual piriform neurons. **A1-4.** Examples from 4 cells showing the relationship between the linearly predicted and observed responses. The size of the symbol indicates the number of odorants in the mixture. Red line shows the fit to equation 1. Dashed line is the unity line. **B.** Histogram of the coefficients of determination (R^2^) of the fits to equation 1 obtained from all cells. **C.** The coefficients of determination plotted against the slope of a linear fit between the observed and linearly predicted responses. **D-F.** Spike width (D), baseline firing rate (E), and preferred respiratory phase (F), plotted against the coefficient of determination obtained in the fit to equation 1.

To assess how the variable fit quality across cells affects population coding in piriform cortex, and how these responses evolve in time, we analyzed response trajectories of 37 neurons that were recorded using the same odorant set (odors 1, 3, 5, and 7). We used principal component analysis to visualize response trajectories (Figure 3). Mixture responses typically occupied the space between the component responses (Figure 3A). To test how well the normalization model explains population responses, we created normalized PSTHs for each mixture and projected them onto PCA space (Figure 3B). Normalized PSTHs were created by simultaneously fitting a single pair of parameters to all mixture responses of each neuron. We tested 3 other models for comparison: 1) a linear model in which the predicted mixture response is the sum of the component responses. 2) a mean model in which the mixture response is predicted to be the average of the component responses. And 3) a max model in which the predicted mixture response is the maximum of the component responses (Figure 3B-C). We compared the errors produced by the four models by measuring the Euclidean distance between the modelled PSTHs and the actual observed responses. Overall the normalization model was the most accurate with the lowest distance for 9 out of the 11 mixtures (Figure 3G-H, p=0.005 vs mean, p=0.001 vs max, p=0.001 vs linear, Wilcoxon signed rank test). To assess the accuracy of the normalization model, we compared its errors to the distances between the 15 different presented stimuli. The model errors were significantly lower than the inter-stimulus distances, despite the fact that our stimuli have many shared odorants (Figure 3I). These results indicate that population coding of mixtures in piriform cortex can be explained by a normalization model.

**Figure 3.**
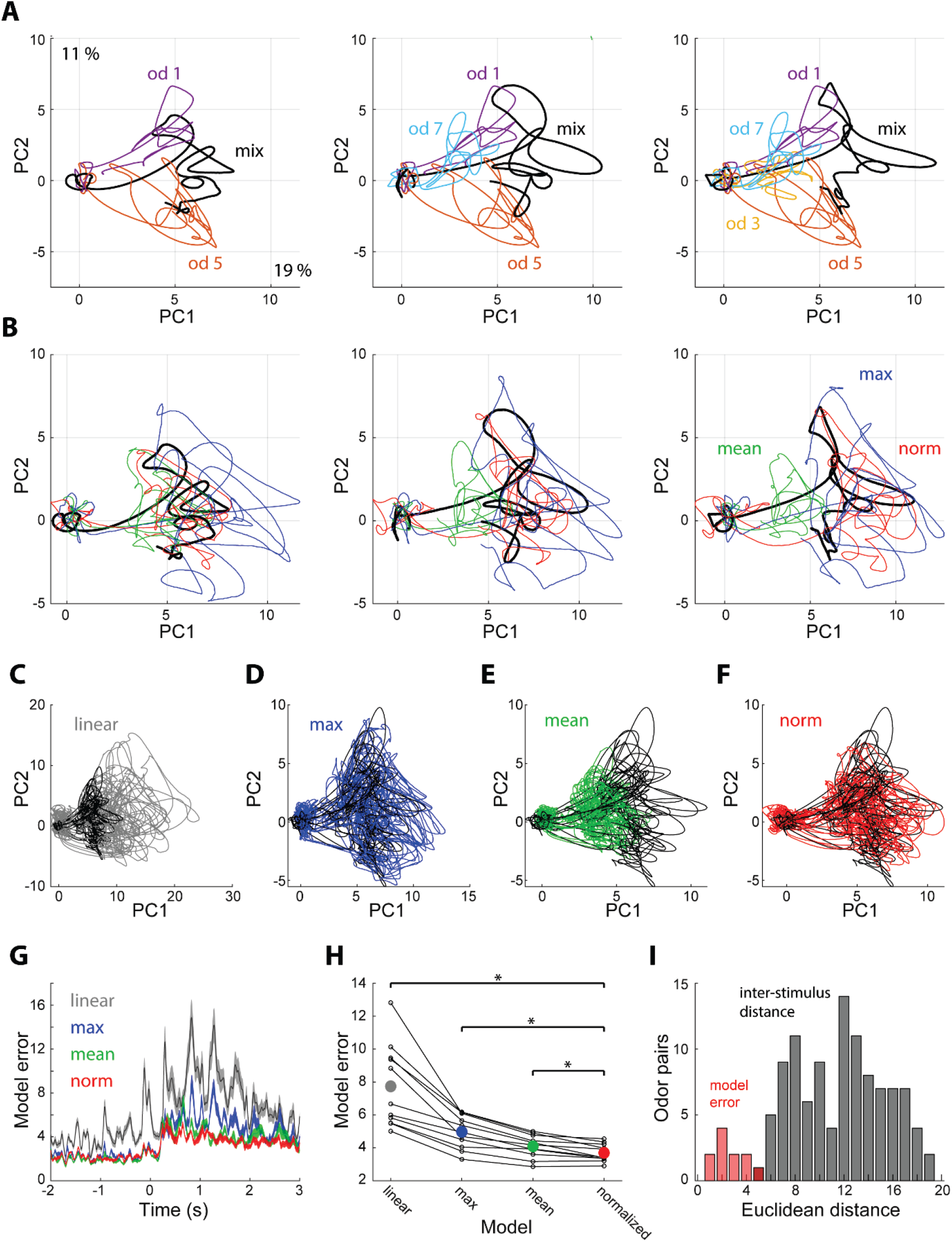
Population coding of mixture responses. **A.** Projections of single odorant (colored) and respective mixture responses (black) onto the first 2 principal components explaining 19 and 11 percent of the variance, respectively. Shown are examples for mixtures of 2,3, and 4 odorants. **B.** PCA projections of the same mixture responses as in A (black) and 3 corresponding predicted responses. Blue – max, Green – mean, Red – normalization. **C-F.** PCA projections of all 11 mixture responses (black) and their corresponding predicted responses (colored). **C.** Linear prediction. **D.** Max prediction. **E.** Mean prediction. **F.** Normalization prediction. **G.** Model errors, calculated as the Euclidean distance between the observed and predicted mixture responses as a function of time. Colors as in B-F. Shown are mean ± SEM. **H.** Mean model error distance during odor presentation for all model-mixture pairs. Dots representing the same mixture are connected with lines. Colored dots show the mean. Asterisks reflect statistical significance (p<0.01). **I.** Normalization model error for the 11 mixtures (red), and all pairwise inter-stimulus distances (gray).

One of the suggested tasks that the piriform cortex is presumed to solve is figure-background segmentation. It was recently suggested that individual piriform neurons may be able to detect target odors in a background-invariant manner by pooling inputs from multiple glomeruli (Mathis et al., 2016; Singh et al., 2019). We therefore asked whether the presence of individual odorants in mixtures can be extracted from the activity of individual piriform neurons. We used receiver operator characteristic (ROC) analysis to test whether the responses to mixtures that contain a specific target odorant are discriminable from responses to mixtures that do not contain the target odorant (Figure 4). For each neuron, we assessed discriminability for all 4 odorants in the experiment and quantified it with the area under the ROC curve (auROC). We assessed each neuron’s discrimination power with its best discriminated odorant (Figures 4A-B). Many neurons yielded high levels of discriminability with 54/56 (96%) neurons performing better than shuffled-labels controls, and 11/56 (20%) cells having an auROC of above 0.8. The discriminability for each odorant based on its single best neuron ranged from auROC of 57 to 99.7 and averaged at 79 (Figure 4C). Mice detect target odorants from background mixtures in about 500 ms (Rokni et al., 2014). We therefore analyzed how single neuron target detection is affected by the time window over which responses are integrated. Responses integrated within a window of less than 200 ms from stimulus onset were not informative about target odorants and the auROC curve plateaued for most neurons between 800 and 1600 ms (Figure 4D). The longer integration windows are presumably more informative about target odorants due to increased spike counts and signal to noise ratio, but they may also be more informative because they cover the right epoch in which action potentials carry the most information about the presence of the target odorant. To analyze for the latter, we repeated the ROC analysis with 200 ms windows that were shifted in time (Figure 4E). 12/56 neurons had at least one 200 ms epoch for which the auROC was above 0.8. The auROC as a function of time for these 12 neurons was highly correlated with the shape of their PSTHs in response to the target odorants. These analyses indicate that single piriform neurons are highly informative about the presence of a target odorant within a mixture, and that detection can be performed in time scales that match previous behavioral measurements.

**Figure 4.**
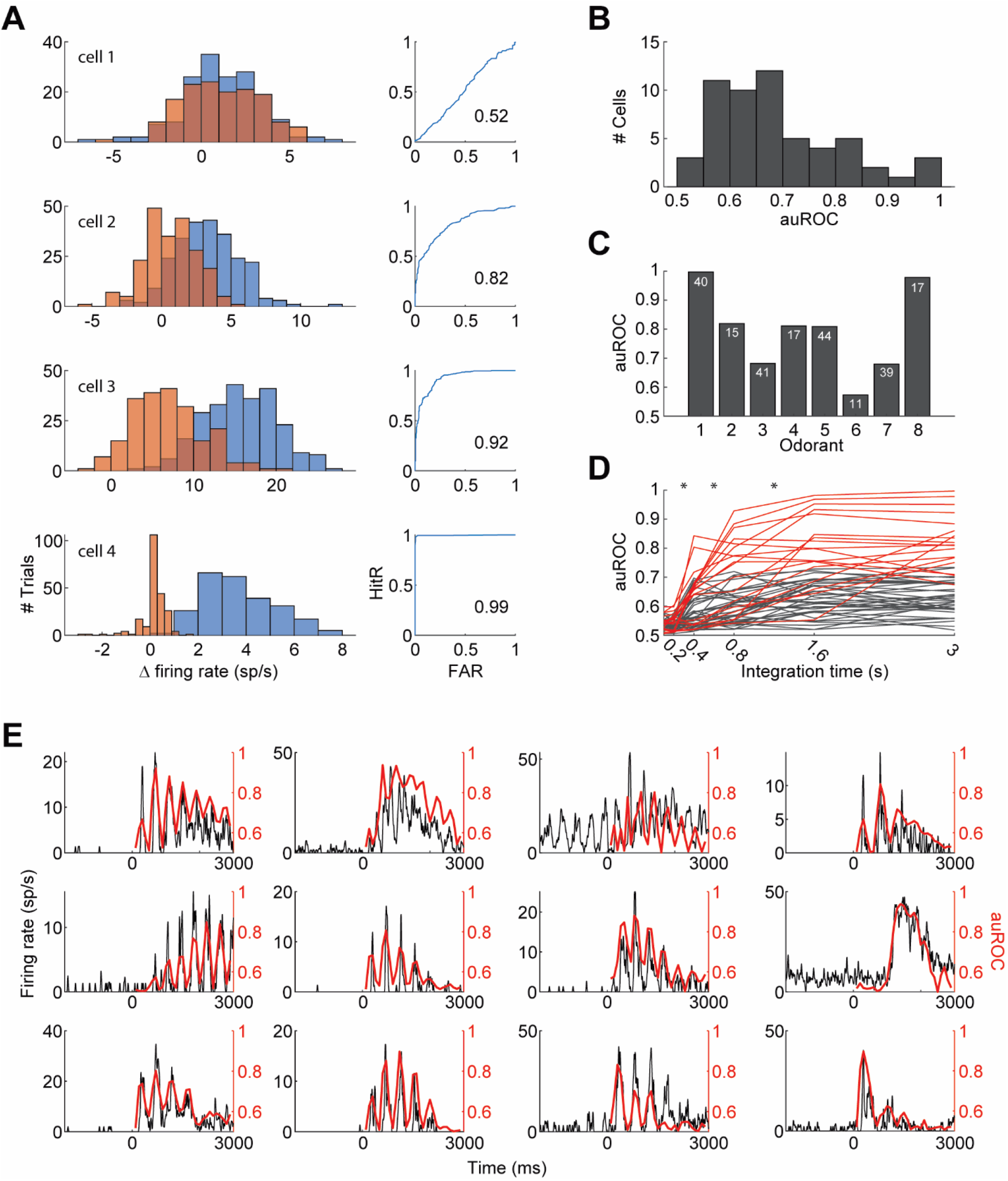
Detection of target odorants with single neurons. **A.** The ROC analysis of 4 example cells. Firing rate histograms are shown on left (blue – all mixtures including a particular odorant, orange – all mixtures excluding the odorant). Panels on the right show hit rate (HitR) vs false alarm rate (FAR) as the decision boundary is shifted (numbers indicate the area under the curve). **B.** Histogram of the area under the ROC curve. The highest value for each neuron (across the 4 odorants) is included. **C.** The maximal area under the curve obtained for each odorant. The number of neurons tested with each odorant is indicated. **D.** The area under the curve for each cell (with its best odor) as a function of the duration of response integration. Low performing cells (never reaching performance of 0.8) are shown in gray, high performing cells are shown in red. **E.** The area under the curve using a sliding 200 ms integration window as a function of the time of integration along the response (red) superimposed with the PSTH in response to the detected odorant (black). 12 high performing neurons are shown.

Despite the overall good performance of target odorant detection form the activity of single neurons, some odorants could not be accurately detected (see odors 3 and 7 in Figure 4C). We therefore next analyzed the ability to detect target odorants from the activity of neuronal populations. We pooled data from all cells stimulated with a specific set of 4 odorants (odorants 1, 3, 5, and 7; 37 cells) to create a pseudo-population. We then trained linear classifiers (realized as logistic regressions) to classify trials according to the presence of each individual odorant. Classifiers were cross-validated by constructing a test set that included a single response from each neuron to each of the 15 odor combinations. Test set responses were not included in the training set. The mean classification accuracy across target odorants was 88 % correct (99% for odorant 1, 88% for odorant 3, 86% for odorant 5, and 78% for odorant 7), showing a significant improvement for the odorants that were not well detected with single neurons. The performance of the classifiers typically decreased for richer mixtures, starting from an average performance of 94±2 % for single odorants and decreasing to 82±6% for mixtures with 4 components (Figure 5A). To assess the number of neurons required to detect individual odorants from mixtures, we trained classifiers using sub-populations (Figure 5B). We started with the entire population and gradually removed the neuron with the minimal absolute weight. On average across odors, 6 neurons were sufficient to reach a performance level above 90 % of the level achieved with the entire population (1 neuron for odorant 1, 8 for odorant 2, 7 for odorant 3, and 6 for odorant 4). We next asked whether target odorants can be detected within behavioral time scales. To that end, we analyzed how classifier performance depends on the response integration time (Figure 5C). Classifier performance degraded only for one target odorant when decreasing the integration time from 3 to 1 s, but degraded sharply for all target odorants when decreasing the integration window below 500 ms. The average performance across odors with 500 ms integration time was 77±5 %, indicating that much of the information for classification is available within the behavioral time limits. Lastly, we tested whether spike timing carries any information beyond what is carried by the mean firing rates. We trained classifiers with temporal inputs at varying temporal resolution (Figure 5D). Decreasing time bins below 500 ms yielded over-fitted models that performed badly on test trials. Even when the total response duration was lowered to 1 second and the number of training trials was increased 100 fold, time bins below 500 ms yielded over-fitted models. This indicates that the anterior piriform cortex uses a rate code to convey information about the presence of target odors embedded in background mixtures. Together, these analyses show that the detection of target odorants against background mixtures can be reliably achieved with a small number of piriform neurons.

**Figure 5.**
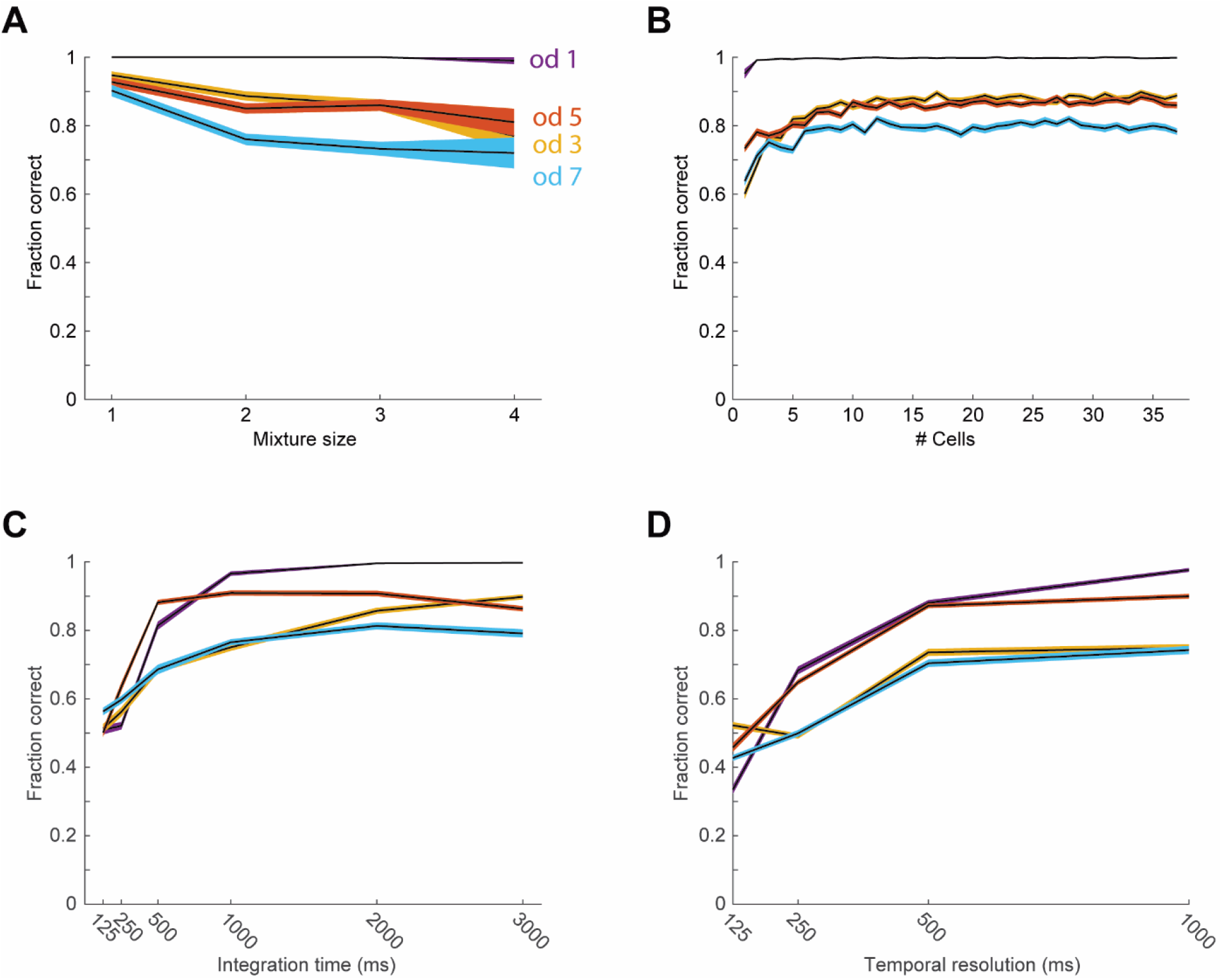
Detection of target odorants with pseudo-populations. In all panels black lines show the average performance for each odorant, and the color-shaded areas show the standard error of the mean. The different colors represent the 4 different odorants (blue – od1, red – od3, yellow – od5, purple – od7). **A.** Performance of the linear classifiers as a function of the number of odorants in the mixture. **B.** Performance of the classifiers as a function of the number of neurons used as input. **C.** Performance of the classifiers as a function of the duration of the response integration window. **D.** Performance of the classifiers as a function of temporal resolution.

## Discussion

We recorded from single neurons in the anterior piriform cortex of naïve anesthetized mice and describe their responses to odorant mixtures. We used receiver operator characteristic (ROC) analysis, and linear classifiers to describe the availability of information regarding mixture components in single neuron, and population activity, respectively. Our main findings are that (i) piriform responses can be described with a normalization model that relates mixture responses to the linear sum of individual component responses, and (ii) individual odorants can be reliably detected from mixtures by the firing of small neuronal populations in the piriform cortex.

Previous analyses of mixture representations in the piriform cortex have found that sub-linear summation of odorant responses is the common case, although examples of supra-linear summation have also been demonstrated (Yoshida and Mori, 2007; Stettler and Axel, 2009). Similar variability in odorant summation has also been found in the anterior olfactory nucleus (Lei et al., 2006). Our analysis shows that combining multiple mixture responses of individual neurons uncovers a mathematical relationship between them. The normalization model may explain some of the variability in previous studies as responses far below the saturation level (R_max_ in eq 1) may appear linear while greater responses will be sublinear.

Normalization has been found in many cortical regions and has been proposed as a canonical computation performed by cortical circuits that allows them to encode variables monotonically without saturating (Carandini and Heeger, 2012). Our finding of mixture response normalization is in line with studies that demonstrated normalization of responses to increasing concentrations of odorants (Bolding and Franks, 2017, 2018; Stern et al., 2018). Several mechanisms may contribute to the sublinear summation of odorant responses. These include interactions between odorants at the olfactory epithelium (Kurahashi et al., 1994; Duchamp-Viret et al., 2003; Oka et al., 2004; Grossman et al., 2008; Takeuchi et al., 2009; Reddy et al., 2018; Xu et al., 2019; Zak et al., 2019), processing in the olfactory bulb (Giraudet et al., 2002; Linster and Cleland, 2004; Tabor et al., 2004; Lin et al., 2006), and further normalization within piriform cortex, implemented by local inhibition. Inhibitory circuits within the piriform cortex have indeed been shown to play a major role in shaping piriform representations (Poo and Isaacson, 2009, 2011; Franks et al., 2011; Bolding and Franks, 2018).

Our findings complement recent studies describing normalization of piriform cortical responses to increasing concentrations (Bolding and Franks, 2017, 2018). Normalization of increasing concentrations is suggested to curtail responses to late activating glomeruli and thereby create concentration invariant representations due to a concentration invariant order of activation of receptors (Wilson et al., 2017; Chong et al., 2020). It is not clear how curtailing responses to late activating glomeruli may serve mixture coding. Adding odorants to a mixture is likely to change the order of activated glomeruli.

Most neurons in our data set were well fit by the normalization model, however, it is important to note that some neurons were not. These cells were mostly cells that did not increase their activity when mixture components were added, and it is possible that we only probed them with stimuli that push them to their saturated response levels. Although the quality of fit to the normalization model was not correlated with any electrophysiological signature (baseline firing, phase preference, and spike waveform), we cannot rule out the possibility that different neuronal populations integrate mixture components differently. Indeed principal neurons in the piriform cortex include subpopulations that differ in their localization across layers, biophysical properties, projection targets, and possibly tuning width (Suzuki and Bekkers, 2006, 2011; Zhan and Luo, 2010; Diodato et al., 2016).

The finding that single piriform neurons may be sufficient to detect target odorants against background mixtures is in line with recently proposed models for target odorant detection from mixtures (Mathis et al., 2016; Singh et al., 2019). In these models, individual glomeruli are not informative about the presence of any specific odorant, but pooling information from a population of glomeruli, supports accurate classification. As piriform neurons receive inputs from multiple receptor channels, they are poised to be suitable for this classification. Importantly however, these studies suggest that the synaptic connections between the olfactory bulb and piriform cortex must be learned for proper classification. Here we show that individual neurons perform this classification in naïve mice. Learning and synaptic plasticity may still improve classification accuracy as suggested by the models. Specific responses to mixture components have also been observed in the insect equivalent of the piriform cortex (but not in the antennal lobe) (Shen et al., 2013). Using linear classifiers, we showed that target odorant detection can be improved beyond what is achieved with single neurons by pooling inputs from a small population of neurons. Providing classifiers with temporal inputs yielded over-fitted models indicating that the anterior piriform cortex probably does not use a temporal code to represent odorants embedded in mixtures. This is in line with previous work indicating that the anterior piriform cortex primarily employs a rate code to project odor information (Miura et al., 2012; Haddad et al., 2013). The inputs for the classifiers were pseudo-populations pooled from multiple experiments. Pooling responses from multiple experiments removes noise-correlations and therefore is expected to improve classifier performance (Zohary et al., 1994). It should be noted however that previous measurements have found very little noise correlations in the piriform cortex during odor sampling (Miura et al., 2012).

In this study we focused on the ability to detect mixture components, however, mixtures of odorants that are emitted from individual objects typically create a unified percept in which the individual components are not identified (Jinks and Laing, 2001; Kadohisa and Wilson, 2006b; Barnes et al., 2008; Gottfried, 2010; Wilson and Sullivan, 2011; Howard and Gottfried, 2014). It will be interesting for future studies to investigate how normalized mixture representations support unified object percepts.

## Acknowledgements

This work was funded by a European Research Council Starting Grant (755764 - COFBMIX). We are grateful to Yoram Ben-Shaul, Mati Joshua, Adi Mizrahi, and Ithai Rabinowitch for commenting on early versions of the manuscript, and to members of the Rokni lab for stimulating discussions. This manuscript has been released as a pre-print at https://www.biorxiv.org/content/10.1101/2019.12.26.888693v4, (Penker et al., 2020)

